# The impact of daily caffeine intake on nighttime sleep: signs of overnight withdrawal?

**DOI:** 10.1101/2020.05.26.114769

**Authors:** Janine Weibel, Yu-Shiuan Lin, Hans-Peter Landolt, Joshua Kistler, Sophia Rehm, Katharina M. Rentsch, Helen Slawik, Stefan Borgwardt, Christian Cajochen, Carolin F. Reichert

**Author notes:** Address for correspondence: Christian Cajochen, PhD, Centre for Chronobiology, Psychiatric Hospital of the University of Basel, Wilhelm Klein-Strasse 27, CH-4002 Basel. Authors contributed equally to this work (Christian Cajochen, Carolin F. Reichert).

## Abstract

Acute caffeine intake can delay sleep initiation and reduce sleep intensity, particularly when consumed in the evening. However, it is not clear whether these sleep disturbances disappear when caffeine is continuously consumed during daytime, which is common for most coffee drinkers. To address this question, we investigated the sleep of twenty male young habitual caffeine consumers during a double-blind, randomized, crossover study including three 10-day conditions: caffeine (3 x 150 mg caffeine daily), withdrawal (3 x 150 mg caffeine for eight days, then switch to placebo), and placebo (3 x placebo daily). After nine days of continuous treatment, electroencephalographically (EEG)-derived sleep structure and intensity were recorded during a scheduled 8-h nighttime sleep episode starting 8 (caffeine condition) and 15 h (withdrawal condition) after the last caffeine intake. Upon scheduled wake up time, subjective sleep quality and caffeine withdrawal symptoms were assessed. Unexpectedly, neither polysomnography-derived total sleep time, sleep latency, sleep architecture, nor subjective sleep quality differed among placebo, caffeine, and withdrawal conditions. Nevertheless, EEG power density in the sigma frequencies (12-16 Hz) during non-rapid eye movement (NREM) sleep was reduced in both caffeine and withdrawal conditions when compared to placebo. These results indicate that daily caffeine intake in the morning and afternoon hours does not strongly impair nighttime sleep structure or subjective sleep quality in healthy good sleepers who regularly consume caffeine. The reduced EEG power density in the sigma range might represent early signs of overnight withdrawal from the continuous presence of the stimulant during the day.

**Statement of Significance:** Caffeine consumption is highly prevalent worldwide and has been repeatedly shown to acutely disrupt sleep, particularly when consumed after several days of abstinence or close to bedtime. However, commonly, caffeine is consumed daily and during daytime. Our well-controlled laboratory study revealed that this common pattern of intake affects nighttime sleep differently: While slow-wave sleep duration or slow-wave activity were rather similar compared to placebo, caffeine intake surprisingly reduced EEG power density in the sigma frequencies. In the light of earlier studies, this might present early signs of caffeine withdrawal which occurs due to overnight caffeine abstinence. The present findings provide novel insights into the impact of daily presence and nightly abstinence of caffeine on nighttime sleep in regular consumers.

## 1. Introduction

Caffeine is the most popular psychoactive substance in the world,^1^ consumed daily by around 80% of the population.^2^ While caffeine is frequently used to counteract sleepiness and boost performance,^3^ its consumption is commonly avoided in the evening^4,5^ to prevent adverse consequences on nocturnal sleep.^3^ The sleep disrupting effects of caffeine are mainly attributed to its influence on the homeostatic component of sleep-wake regulation. Sleep homeostasis describes the increase in sleep pressure during time awake and its dissipation during the following sleep episode,^6^ which has been suggested to be related to rising and decreasing concentrations of adenosine.^7^ Caffeine is an adenosine receptor antagonist, which blocks the A_1_ and A_2A_ adenosine receptors in the central nervous system.^1^ It may, thus, attenuate the increase in sleep pressure during wakefulness^8^ and lead to delayed sleep initiation and more superficial sleep.^9^

The effects of caffeine intake on the quality and quantity of sleep depend on the timing of its consumption. More specifically, caffeine consumed in the evening hours prolongs sleep latency,^10-14^ reduces total sleep time (TST),^10-12,14,15^ shortens deep sleep,^10,12-15^ and decreases electroencephalographically (EEG)-derived slow-wave activity (SWA),^10^ while activity in the sigma range is increased.^10^ However, evening caffeine intake only accounts for approximately 10-20% of the total daily caffeine intake in regular consumers.^4,5^ It needs to be elucidated whether habitual caffeine intake restricted to the morning and afternoon hours similarly affects nighttime sleep.

Furthermore, not only the timing but also the frequency of preceding caffeine intake prior to sleep may be an important factor for the repercussions on sleep. The majority of the worldwide population consumes caffeine on a daily basis,^2^ which can lead to tolerance development due to the recurrent supply of the psychostimulant.^1^ In line with these results, the sleep-disrupting effects of continuous high-dose caffeine in the morning, afternoon, and evening (3 x 400 mg) intake vanished and only stage 4 sleep remained reduced after one week of caffeine intake.^12^ However, whether more sensitive markers for sleep intensity such as spectral sleep EEG measures, adapt to the long-term exposure to the stimulant has to our best knowledge not yet been investigated.

Importantly, not only caffeine *per se*, but also the state of acute abstinence to which regular consumers expose themselves every night, might affect sleep. This so-called overnight abstinence represents the start of a caffeine withdrawal phase.^16^ Withdrawal symptoms such as increased tiredness,^17^ longer sleep duration and better sleep quality^18^ can be observed at a subjective level starting roughly 12 h after last caffeine intake.^17^ However, the influence of caffeine withdrawal on objective EEG-derived sleep variables were not systematically reported up to date and remain to be compared against a placebo-baseline.

Here we aimed at determining whether daily caffeine intake during morning and afternoon hours impairs nighttime sleep structure and sleep intensity after continuous daytime caffeine intake over nine days. We hypothesized a reduced depth of sleep after caffeine intake, indexed in shortened slow-wave sleep (SWS) duration and a decrease in SWA compared to placebo. Moreover, we hypothesized that the abrupt cessation from the daily intake generates acute subjective withdrawal symptoms, and changes sleep structure and intensity compared to both the daily caffeine intake and the placebo-baseline.

## 2. Methods

### 2.1 Participants

Twenty male volunteers were recruited into the present study through online advertisements and flyers distributed in public areas. Interested individuals aged between 18 and 32 years old (mean age ± SD: 26.4 ± 4 years) and reporting a daily caffeine consumption between 300 and 600 mg (mean intake ± SD: 478.1 ± 102.8 mg) were included. The self-rating assessment for the daily amount of caffeine intake was structured based on Bühler et al.,^19^ and the amount of caffeine content was defined according to Snel and Lorist.^3^ To ensure good health, volunteers were screened by self-report questionnaires and a medical examination conducted by a physician. Additionally, all volunteers reported good sleep quality assessed with the Pittsburgh Sleep Quality Index (PSQI; score ≤ 5)^20^ and showed no signs of sleep disturbances in a polysomnography (PSG) recorded during an adaptation night in the laboratory (sleep efficiency (SE) > 70%, periodic leg movements < 15/h, apnea index < 10). To control for circadian misalignment, volunteers who reported shiftwork within three months and transmeridian travels (crossing > 2 time zones) within one month prior to study admission were excluded. Further exclusion criteria comprised body mass index (BMI) < 18 or > 26, smoking, drug use, and extreme chronotype assessed by the Morningness-Eveningness Questionnaire (MEQ; score ≤ 30 and ≥ 70).^21^ To reduce variance in the data incurred by the effect of menstrual cycle on sleep^22^ and the interaction between caffeine metabolism and the use of oral contraceptives,^23,24^ only male volunteers were studied. A detailed description of the study sample can be found in Weibel et al.^25^

All volunteers signed a written informed consent and received financial compensation for study participation. The study was approved by the local Ethics Committee (EKNZ) and conducted according to the Declaration of Helsinki.

### 2.2 Design and Protocol

We employed a double-blind randomized crossover study including a caffeine, a withdrawal, and a placebo condition. Volunteers were allocated to the order of the three conditions based on pseudo-randomization, for more details see Weibel et al.^25^ As illustrated in Figure 1, each condition started with an ambulatory part of nine days, followed by a laboratory part of 43 h. In each condition, participants took either caffeine (150 mg) or placebo (mannitol) in identical appearing gelatin capsules (Hänseler AG, Herisau, Switzerland) three times daily, scheduled at 45 min, 255 min, and 475 min after awakening, for a duration of 10 days. This regimen was applied based on a previous study investigating tolerance to the effects of caffeine and caffeine cessation.^18^ To enhance caffeine withdrawal in the withdrawal condition, treatment was abruptly switched from caffeine to placebo on day nine of the protocol (255 min after wake-up, 15 h before sleep recording).

**Figure 1.**
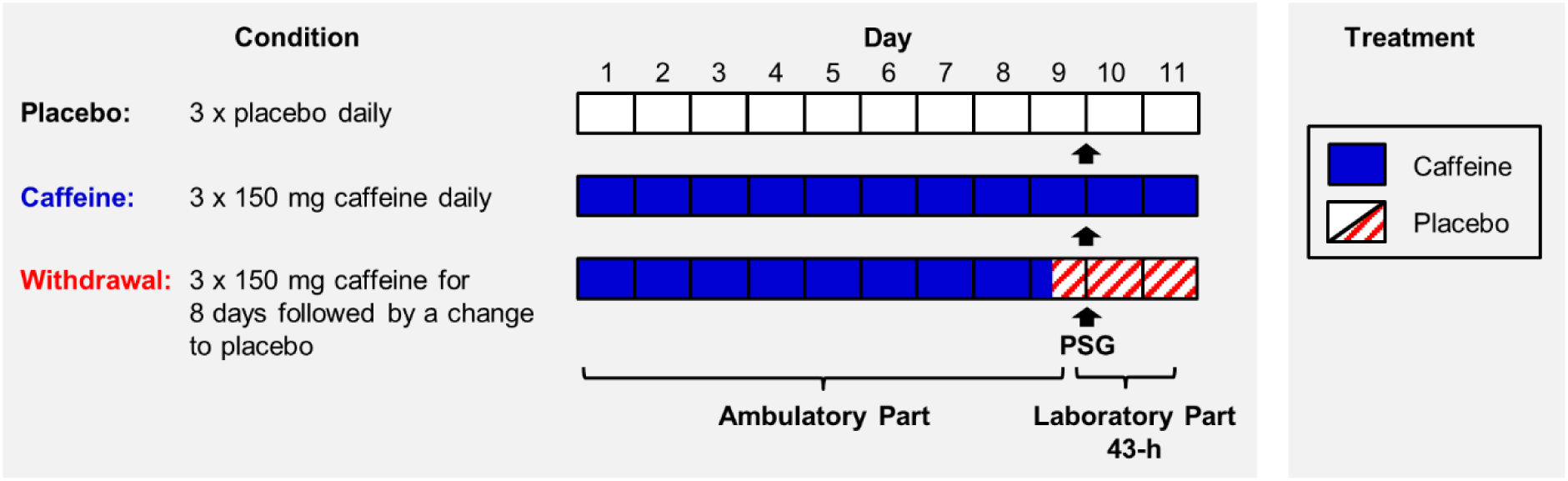
Illustration of the study design. Twenty volunteers participated in a placebo, a caffeine, and a withdrawal condition during which they ingested either caffeine or placebo capsules three times daily (wake-up +45 min, +255 min, and +475 min). Each condition started with an ambulatory part of nine days and was followed by a laboratory part of 43 h. After nine days of continuous treatment, we recorded 8 h of polysomnography (PSG), indicated as arrows, during nighttime sleep under controlled laboratory conditions. The sleep episode was scheduled to volunteers’ habitual bedtime.

During the nine days of the ambulatory part, volunteers were asked to maintain a regular sleep-wake rhythm (± 30 min of self-selected bedtime/wake-up time, 8 h in bed, no daytime napping), verified by wrist actimetry (Actiwatch, Cambridge Neurotechnology Ltd., Cambridge, United Kingdom), and to keep subjective sleep logs. The duration of the ambulatory part was set for nine days based on the maximum duration of withdrawal symptoms^17^ and thus, to avoid carry-over effects from the previous condition. Furthermore, volunteers were requested to refrain from caffeinated beverages and food (e.g. coffee, tea, soda drinks, and chocolate), alcohol, nicotine, and medications. Caffeine abstinence and compliance to the treatment requirements were checked by caffeine levels from the daily collection of fingertip sweat.

On day nine, volunteers admitted to the laboratory at 5.5 h prior to habitual sleep time. Upon arrival, a urinary drug screen (AccuBioTech Co., Ltd., Beijing, China) was performed to ensure drug abstinence. Electrodes for the PSG were fitted and salivary caffeine levels collected. An 8-h nighttime sleep episode was scheduled at volunteers’ habitual bedtime starting 8 and 15 h after the last caffeine intake in the caffeine and withdrawal condition, respectively. The next day, volunteers rated their subjective sleep quality by the Leeds Sleep Evaluation Questionnaire (LSEQ)^26^ and potential withdrawal symptoms by the Caffeine Withdrawal Symptom Questionnaire (CWSQ).^27^

To reduce potential masking effects on our outcome variables, volunteers were housed in single apartments under dim-light (< 8 lx) during scheduled wakefulness and 0 lx during sleep. Volunteers were asked to maintain a semi-recumbent position during wakefulness, except for restroom breaks. Social interactions were restricted to team members and no time-of-day cues were provided throughout the in-lab protocol.

### 2.3 Salivary Caffeine

To characterize individual caffeine levels during nighttime sleep, we report salivary caffeine levels assessed 3 h prior to the scheduled sleep episode and 5 min after wake-up. Samples were stored at 5°C following collection, later centrifuged (3000 rpm for 10 min) and subsequently kept at -28°C until analyses. Liquid chromatography coupled to tandem mass spectrometry was used to analyze the levels of caffeine. One dataset in the withdrawal condition was lost.

### 2.4 Subjective Sleep Quality

Subjective sleep quality was assessed 10 min upon scheduled wake-up time with a paper and pencil version of the LSEQ.^26^ Volunteers were asked to rate 10 items on visual analogue scales which are grouped into four domains (getting to sleep (GTS), quality of sleep (QOS), awake following sleep (AFS), and behavior following wakening (BFW)).

### 2.5 Polysomnographic Recordings

PSG was continuously recorded during 8 h of nighttime sleep using the portable V-Amp device (Brain Products GmbH, Gilching, Germany). Grass gold cup electrodes were applied according to the international 10-20 system including two electrooculographic, two electromyographic, two electrocardiographic, and six electroencephalographic derivations (F3, F4, C3, C4, O1, O2). Channels were referenced online against the linked mastoids (A1, A2). Signals were recorded with a sampling rate of 500 Hz and a notch filter was online applied at 50 Hz.

Each epoch of 30 sec of the recorded PSG data was visually scored according to standard criteria^28^ by three trained team members blind to the condition. SWS was additionally classified into stage 3 and 4 based on Rechtschaffen and Kales.^29^ The scoring agreement between the three scorers was regularly confirmed to reach > 85%.

TST was defined as the sum of the time spent in sleep stages 1-4 and rapid eye movement sleep (REM). Sleep latencies were calculated as minutes to the first occurrence of the corresponding sleep stage following lights off. Non-rapid eye movement (NREM) sleep was calculated as sum of sleep stages 2, 3 and 4. All sleep stages are expressed as relative values (%) of TST.

Spectral analysis was performed by applying fast Fourier transformation (FFT; hamming, 0% overlapped, 0.25 Hz bins) on 4-s time windows. Artifacts were manually removed based on visual inspection, and data were log-transformed prior to spectral analyses. All-night EEG power density during NREM sleep was analyzed for each 0.25 Hz frequency bin in the range of 0.75-32 Hz recorded over the central derivations (C3, C4). SWA was defined as EEG power density between 0.75-4.5 Hz and sigma activity between 12-16 Hz. Sleep cycles were defined based on adapted rules developed by Feinberg and Floyd^30^ and divided into 10 NREM and four REM intervals within each cycle. Ten nights were excluded from sleep analyses due to technical problems (placebo: *n* = 3; caffeine: *n* = 4; withdrawal: *n* = 3).

### 2.6 Caffeine Withdrawal Symptoms

Withdrawal symptoms were first assessed 35 min after wake-up and subsequently prior to each treatment administration with the self-rating CWSQ.^27^ Twenty-three items are grouped into seven factors (fatigue/drowsiness, low alertness/difficulty concentrating, mood disturbances, low sociability/motivation to work, nausea/upset stomach, flu-like feelings, headache) and were rated on a 5 point scale by choosing between 1 (not at all) and 5 (extremely). Prior to analyses, eight items have been reversed scored as they were positively worded (e.g. alert or talkative) in the questionnaire. To assess caffeine withdrawal, we first calculated a sum score comprising all 23 items of the caffeine withdrawal questionnaire. Missing responses to single items were replaced by the median response of each condition over all volunteers in the respective time of assessment. In a next step, we calculated relative withdrawal symptoms in the caffeine and withdrawal condition (i.e. the difference of the withdrawal score in the caffeine and withdrawal condition respectively minus the score of the placebo condition). The data of one volunteer was lost due to technical difficulties.

### 2.7 Statistical Analyses

Analyses were performed with the statistical package SAS (version 9.4, SAS Institute, Cary, NC, USA) by applying mixed model analyses of variance for repeated measures (PROC MIXED) with the repeated factors ‘condition’ (placebo, caffeine, withdrawal) and/or ‘time’ (levels differ per variable) and the random factor ‘subject’. The LSMEANS statement was used to calculate contrasts and degrees of freedom were based on the approximation by Kenward and Roger.^31^ Post-hoc comparisons were adjusted for multiple comparisons by applying the Tukey-Kramer method. A statistical significance was defined as *p* < 0.05. One dataset has been excluded from all the analyses due to non-compliance with the treatment requirements (caffeine: *n* = 1).

## 3. Results

### 3.1 Salivary Caffeine Levels

Caffeine levels significantly differed between each of the three conditions (main effect of condition: *F*_2,90.7_ = 46.12; *p* < 0.001) with the highest levels in the caffeine condition and the lowest in the placebo condition (post-hoc comparisons: *p*_all_ < 0.01). In addition, a significant interaction of the factors condition and time (*F*_2,89.6_ = 10.65; *p* < 0.001) confirmed that caffeine levels were modulated by time with levels decreasing during nighttime sleep in the caffeine condition only (post-hoc comparison: *p* < 0.001), see Figure 2.

**Figure 2.**
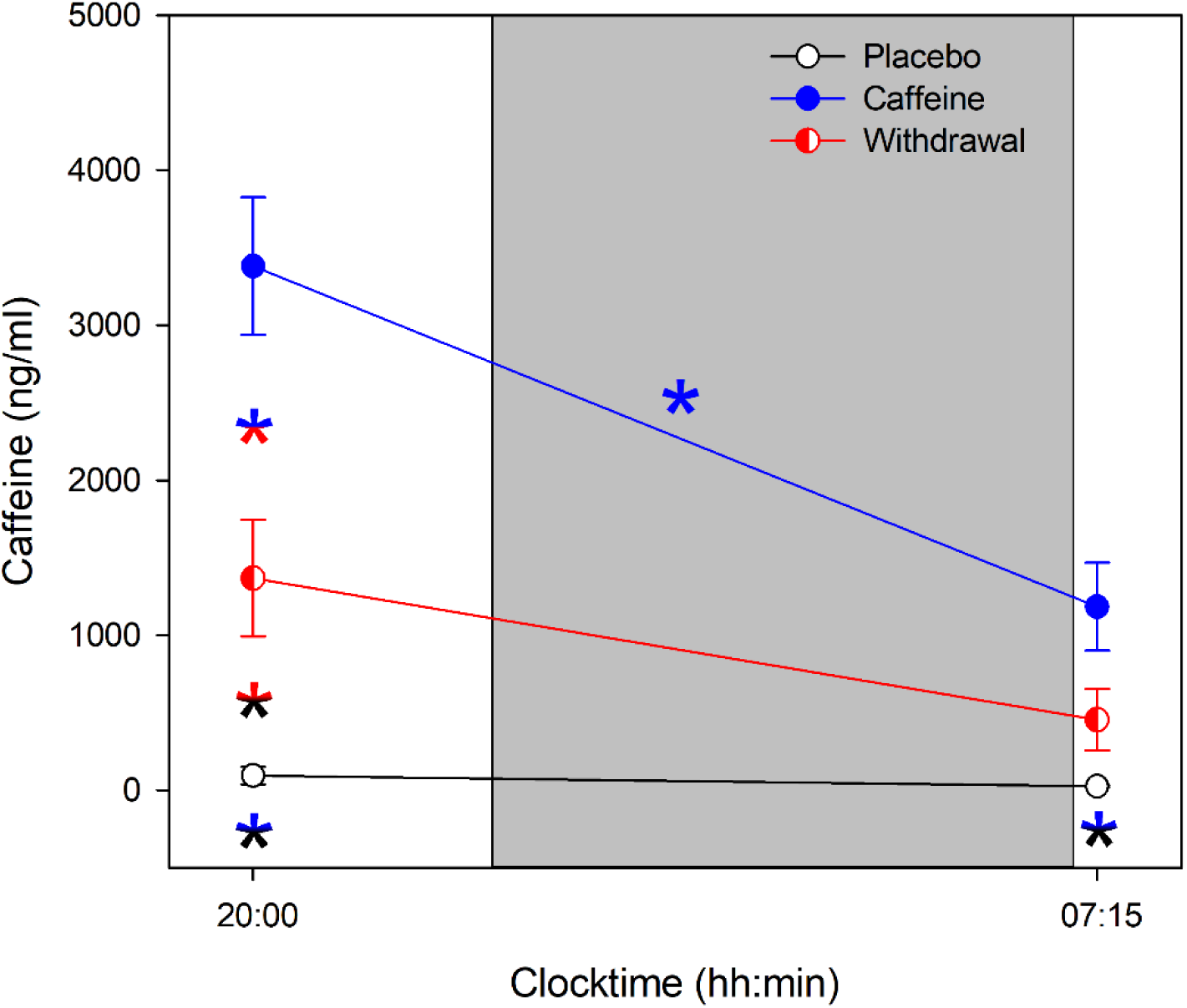
Average caffeine levels collected prior to and after nighttime sleep (grey bar) in the placebo (black open circles), caffeine (blue filled circles), and withdrawal (red semi-filled circles) condition (mean values ± standard errors). The x-axis indicates the mean time of day of sample collection and color-coded asterisks represent significant (*p* < 0.05) post-hoc comparisons of the interaction effect condition x time.

### 3.2 Sleep

Table 1 summarizes the statistical analyses of subjective sleep quality and objective sleep structure assessed during nighttime sleep. Analyses of subjective sleep quality assessed with the LSEQ questionnaire did not reveal significant differences among the three conditions in any of the four domains of sleep quality (*p*_all_ > 0.05).

**Table 1.**
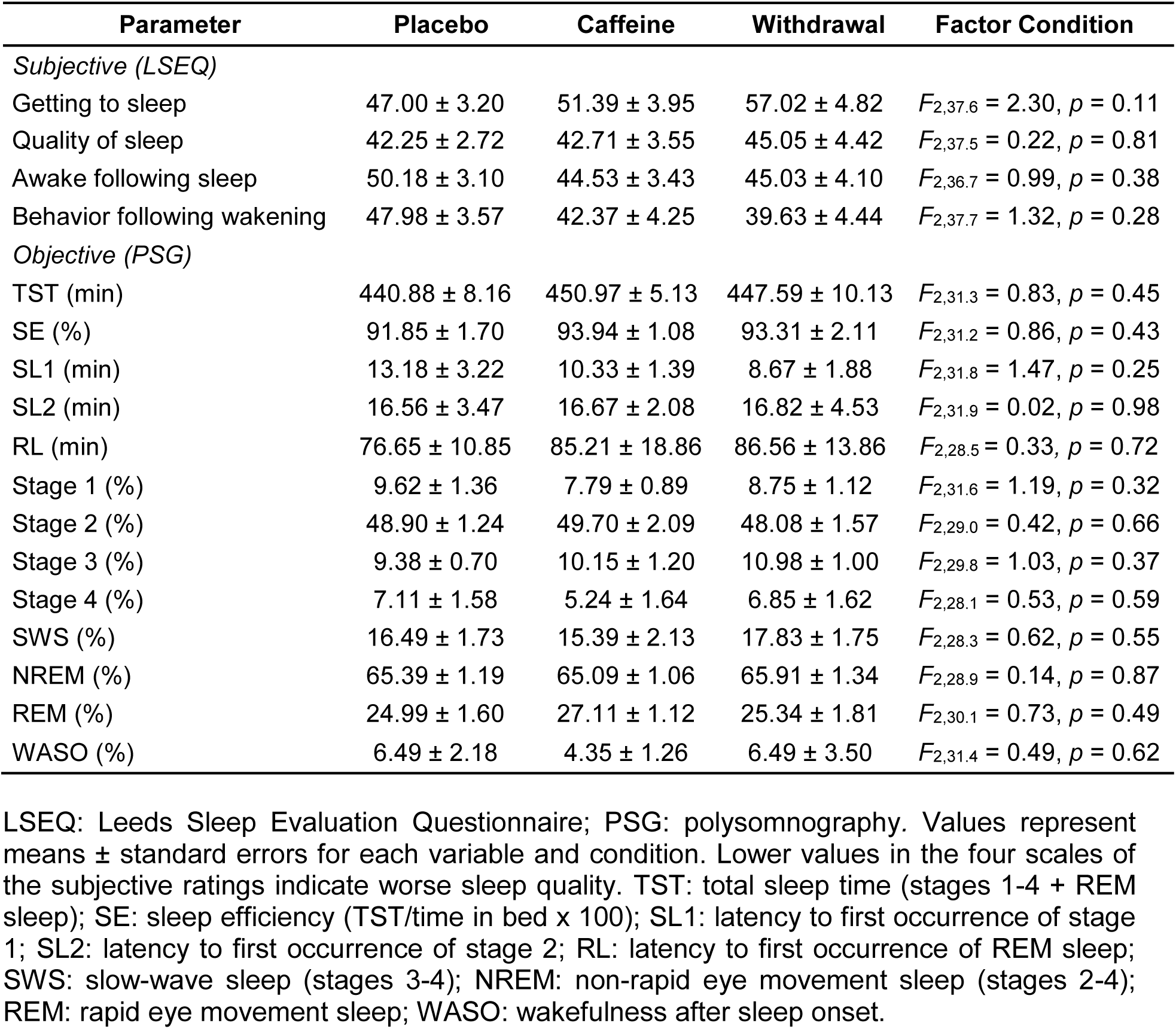
Subjective and objective sleep parameters per condition and results of the analyses.

| Parameter | Placebo | Caffeine | Withdrawal | Factor Condition |
| --- | --- | --- | --- | --- |
| *Subjective (LSEQ)* |  |  |  |  |
| Getting to sleep | 47.00 ± 3.20 | 51.39 ± 3.95 | 57.02 ± 4.82 | $F_{2,37.6} = 2.30, p = 0.11$ |
| Quality of sleep | 42.25 ± 2.72 | 42.71 ± 3.55 | 45.05 ± 4.42 | $F_{2,37.5} = 0.22, p = 0.81$ |
| Awake following sleep | 50.18 ± 3.10 | 44.53 ± 3.43 | 45.03 ± 4.10 | $F_{2,36.7} = 0.99, p = 0.38$ |
| Behavior following wakening | 47.98 ± 3.57 | 42.37 ± 4.25 | 39.63 ± 4.44 | $F_{2,37.7} = 1.32, p = 0.28$ |
| *Objective (PSG)* |  |  |  |  |
| TST (min) | 440.88 ± 8.16 | 450.97 ± 5.13 | 447.59 ± 10.13 | $F_{2,31.3} = 0.83, p = 0.45$ |
| SE (%) | 91.85 ± 1.70 | 93.94 ± 1.08 | 93.31 ± 2.11 | $F_{2,31.2} = 0.86, p = 0.43$ |
| SL1 (min) | 13.18 ± 3.22 | 10.33 ± 1.39 | 8.67 ± 1.88 | $F_{2,31.8} = 1.47, p = 0.25$ |
| SL2 (min) | 16.56 ± 3.47 | 16.67 ± 2.08 | 16.82 ± 4.53 | $F_{2,31.9} = 0.02, p = 0.98$ |
| RL (min) | 76.65 ± 10.85 | 85.21 ± 18.86 | 86.56 ± 13.86 | $F_{2,28.5} = 0.33, p = 0.72$ |
| Stage 1 (%) | 9.62 ± 1.36 | 7.79 ± 0.89 | 8.75 ± 1.12 | $F_{2,31.6} = 1.19, p = 0.32$ |
| Stage 2 (%) | 48.90 ± 1.24 | 49.70 ± 2.09 | 48.08 ± 1.57 | $F_{2,29.0} = 0.42, p = 0.66$ |
| Stage 3 (%) | 9.38 ± 0.70 | 10.15 ± 1.20 | 10.98 ± 1.00 | $F_{2,29.8} = 1.03, p = 0.37$ |
| Stage 4 (%) | 7.11 ± 1.58 | 5.24 ± 1.64 | 6.85 ± 1.62 | $F_{2,28.1} = 0.53, p = 0.59$ |
| SWS (%) | 16.49 ± 1.73 | 15.39 ± 2.13 | 17.83 ± 1.75 | $F_{2,28.3} = 0.62, p = 0.55$ |
| NREM (%) | 65.39 ± 1.19 | 65.09 ± 1.06 | 65.91 ± 1.34 | $F_{2,28.9} = 0.14, p = 0.87$ |
| REM (%) | 24.99 ± 1.60 | 27.11 ± 1.12 | 25.34 ± 1.81 | $F_{2,30.1} = 0.73, p = 0.49$ |
| WASO (%) | 6.49 ± 2.18 | 4.35 ± 1.26 | 6.49 ± 3.50 | $F_{2,31.4} = 0.49, p = 0.62$ |

LSEQ: Leeds Sleep Evaluation Questionnaire; PSG: polysomnography. Values represent means ± standard errors for each variable and condition. Lower values in the four scales of the subjective ratings indicate worse sleep quality. TST: total sleep time (stages 1-4 + REM sleep); SE: sleep efficiency (TST/time in bed x 100); SL1: latency to first occurrence of stage 1; SL2: latency to first occurrence of stage 2; RL: latency to first occurrence of REM sleep; SWS: slow-wave sleep (stages 3-4); NREM: non-rapid eye movement sleep (stages 2-4); REM: rapid eye movement sleep; WASO: wakefulness after sleep onset.

In line with these results, the analyses of the PSG did not reveal significant differences in total sleep time (TST), sleep efficiency (SE), sleep latencies (SL), or the relative amount of sleep stages among the three conditions (*p*_all_ > 0.05).

In a next step, we analyzed all-night EEG power density in the range of 0.75-32 Hz over the central derivations recorded during NREM sleep. In contrast to our assumptions, we did not find any significant differences among the three conditions in the lower frequency bins (0.75-13.25 Hz; *p*_all_ > 0.05). However, power density was significantly reduced compared to placebo in the sigma range during both withdrawal (frequency bins 13.5-17.25 Hz and 18-18.5 Hz, *p*_all_ < 0.05) and caffeine (frequency bins 13.5-16 Hz, *p*_all_ < 0.05).

In a second step, we were interested in the temporal dynamics of both SWA and sigma activity across the night assessed during NREM sleep. As depicted in Figure 3 (top panel), SWA showed a typical temporal pattern with increased activity during the first NREM cycle followed by a steady decline across the night (main effect of time: *F*_39,613_ = 26.28, *p* < 0.001). However, differences among the three conditions did not reach significance (main effect of condition: *F*_2,178_ = 1.33, *p* = 0.27). Also, the interaction of condition and time was not significant (*F*_78,1060_ = 0.89, *p* = 0.74).

**Figure 3.**
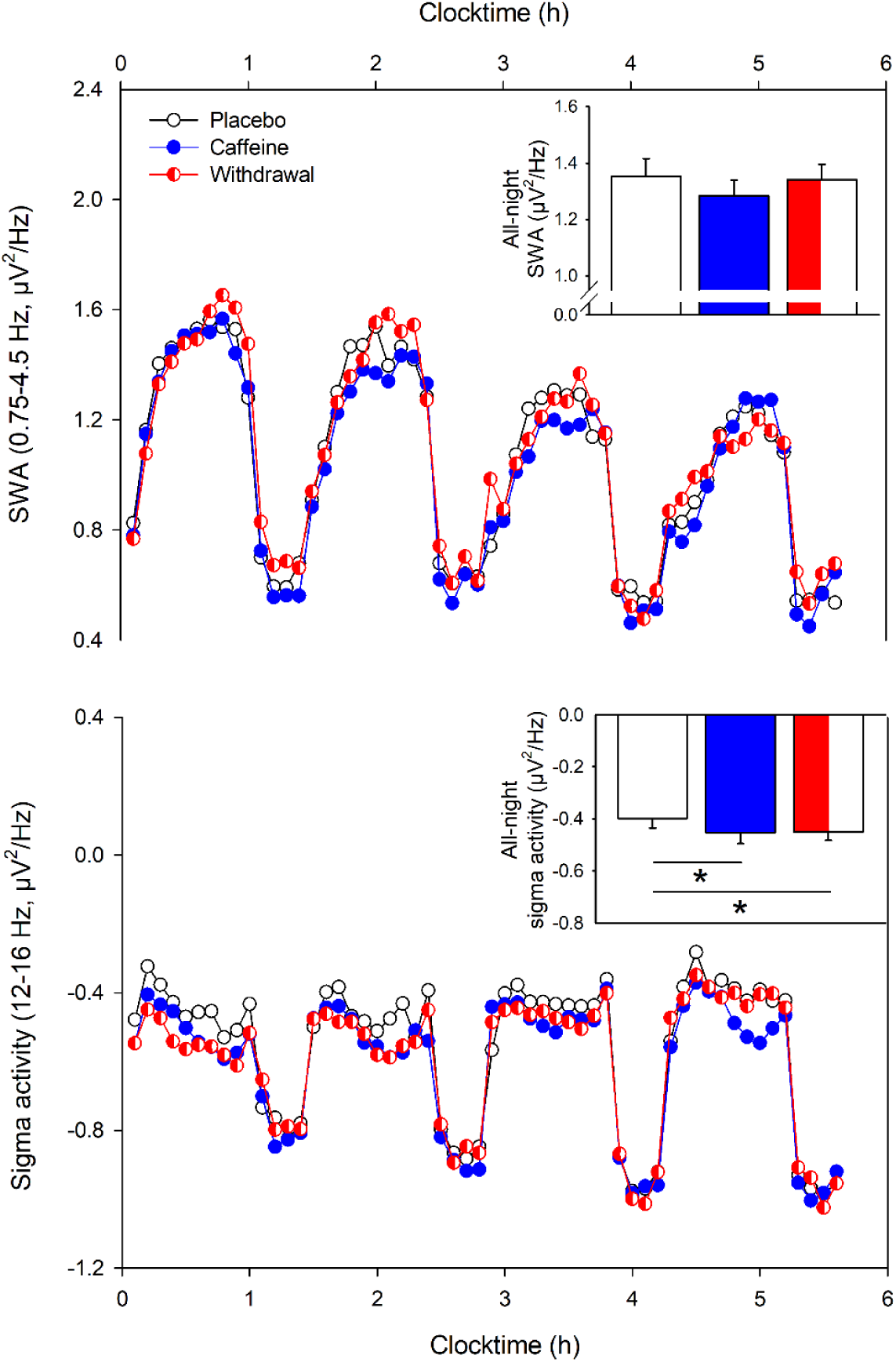
Temporal dynamics of SWA (top) and sigma activity (bottom) during the first four sleep cycles in the placebo (black open circles), caffeine (blue filled circles), and the withdrawal (red semi-filled circles) condition (mean values). The x-axis indicates the mean time of day. While SWA (0.75-4.5 Hz) was not significantly affected by the treatment, sigma activity (12-16 Hz) showed reduced activity during both caffeine and withdrawal compared to the placebo condition (*p*_all_ < 0.05). The inset in each right upper corner represents the mean values ± standard errors of the all-night SWA and sigma activity respectively during NREM sleep in the placebo, caffeine, and withdrawal condition. While all-night SWA (0.75-4.5 Hz) did not differ among the conditions, sigma activity (12-16 Hz) was lower in the caffeine and withdrawal condition compared to placebo (*p* < 0.05). All analyses are based on log-transformed data.

As illustrated in Figure 3 (bottom panel), sigma activity was significantly reduced in both the caffeine and withdrawal conditions compared to placebo intake (main effect of condition: *F*_2,209_ = 19.96, *p* < 0.001; post-hoc comparisons: *p* < 0.001) and the interaction of condition and time tended to be significant (*F* _78,1049_ = 1.25, *p* = 0.08).

Taken together, we could not confirm our assumption of a caffeine-induced reduction of sleep depth, neither in terms of shorter SWS nor in terms of reduced SWA in the caffeine compared to the placebo condition. Based on the discrepancies between the present results and a previous study about the effects of chronic caffeine intake on sleep,^12^ we thus explored whether differences in the individual levels of caffeine before sleep could explain the variance within SWS and SWA. However, no significant effects were observed when controlling for dependent observations within subjects (*p* > 0.05).

### 3.3 Subjective Caffeine Withdrawal Symptoms

Analyses of the relative withdrawal symptoms yielded a significant main effect of condition (*F*_2,20.2_ = 11.30, *p* < 0.01) indicating more withdrawal symptoms during the withdrawal compared to the caffeine condition (post-hoc comparison: *p* < 0.01), depicted in Figure 4. This effect was modulated by time (interaction of condition x time: *F*_2,37.2_ = 3.43, *p* = 0.04), such that the increase in symptoms during the withdrawal compared to caffeine condition was particularly present during the last measurement (*p* < 0.01), i.e. 31 h after the last caffeine intake in the withdrawal condition.

**Figure 4.**
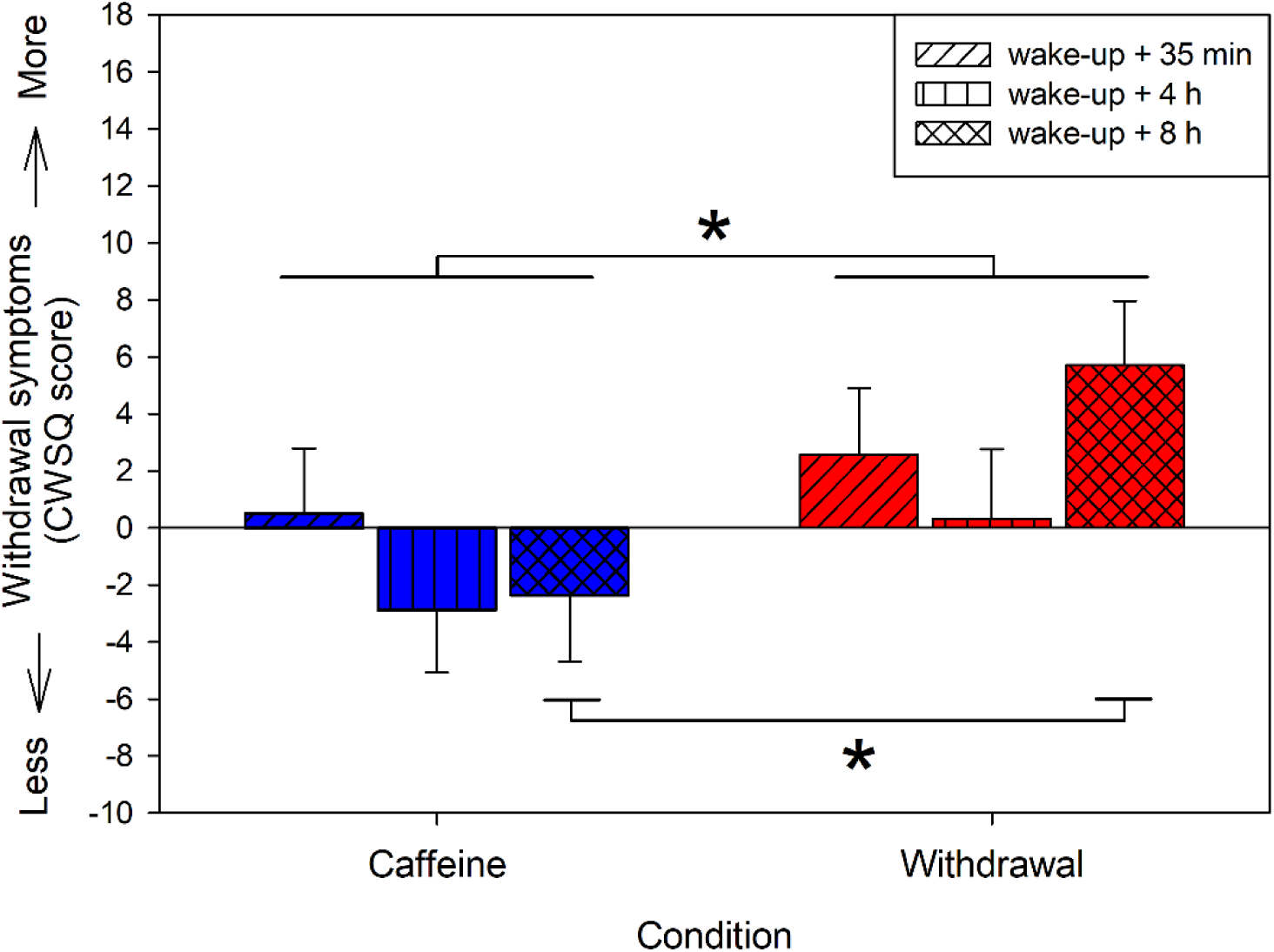
Relative withdrawal symptoms in the caffeine and withdrawal condition (i.e. withdrawal score of the caffeine and withdrawal condition respectively minus the score of the placebo condition) assessed 35 min, 4 h, and 8 h after wake-up on day ten of treatment. Depicted are mean values and standard errors of the relative values (i.e. difference to placebo). Overall, volunteers reported more withdrawal symptoms in the withdrawal condition compared to the caffeine condition (*p* < 0.05). This difference was particularly present 8 h after wake-up during withdrawal compared to caffeine (*p* < 0.001).

## 4. Discussion

The aim of the present study was to investigate the influence of daily daytime caffeine intake and its cessation on nighttime sleep in habitual caffeine consumers under strictly controlled laboratory conditions. Strikingly, caffeine consumption did not lead to clear-cut changes in nighttime sleep structure nor in subjective sleep quality when assessed 8 and 15 h after the last intake in the caffeine and withdrawal condition, respectively. The evolution of subjective withdrawal symptoms indicates that withdrawal becomes perceivable at earliest between 27-31 h after intake. However, compared to placebo, EEG power density was reduced in the sigma range during both caffeine and withdrawal conditions. We conclude that daily daytime intake of caffeine does not strongly influence nighttime sleep structure nor subjective sleep quality in healthy men when consumed in the morning, midday, and in the afternoon. In contrast to the reported increases in sigma activity after acute caffeine intake,^10^ the observed changes in the sigma frequencies might point to early signs of caffeine withdrawal which occur due to overnight abstinence and presumably derive from preceding caffeine-induced changes in adenosine signaling.

To quantify the influence of caffeine on sleep, the stimulant is commonly administered close to the onset of a sleep episode,^10-14^ for instance within one hour prior to bedtime.^10,11,13,14^ Taking into account that caffeine plasma levels peak within 30 to 75 min following caffeine ingestion,^32^ consumption within one hour prior to sleep allows the stimulant to exert its maximum effects at sleep commencement. Indeed, the sleep disrupting effects of caffeine are frequently reported to affect sleep initiation or the first half of the sleep episode.^10-14^ Moreover, sleep intensity, which is usually strongest at the beginning of the night,^33^ was particularly disrupted during the first sleep cycle, as indexed in reduced SWS and SWA.^10^ However, caffeine intake in the evening, particularly after 9 pm is rare,^5^ presumably to avoid impairment of subsequent sleep.^3^ Up to date it remained fairly unclear whether caffeine intake in the morning and afternoon still bears the potential to disrupt nighttime sleep. Thus, our data provide first evidence that daily daytime caffeine intake does not necessarily alter subsequent sleep structure and SWA when consumed > 8 h prior to sleep. Importantly, our findings do not preclude potential impairments of nighttime sleep after morning caffeine intake, if preceded by several days of abstinence from the stimulant.^34^ It rather appears likely that the duration of preceding caffeine consumption drives the discrepancies between acute and chronic effects of caffeine on sleep.

Chronic caffeine intake induces some tolerance development in both physiological measures such as cortisol,^35^ blood pressure,^36^ heart rate,^37^ and also subjective measures such as alertness.^18^ Over time, the stimulatory effects of the substance vanish potentially due to changes in adenosine levels^38^ and/or adenosine receptors.^39-42^ Accordingly, an one-week treatment of caffeine reduced the sleep disrupting effects on deep sleep, even under conditions of high evening dosages.^12^ Thus, the available evidence and the absence of clear-cut changes in the present study point to adaptive processes in sleep initiation, sleep structure, and subjective sleep quality due to the long-term exposure to the stimulant.

However, chronic caffeine consumption bears the risk of withdrawal symptoms when abruptly ceased. These symptoms have been reported to occur as early as 6 h but with peak intensity being reached within 24 to 51 h after last caffeine intake.^17^ While 25 h of caffeine abstinence might not affect nighttime sleep structure,^12^ 32 h of abstinence improved subjective sleep quality.^18^ Thus, scheduling the start of the sleep episode to 15 h after the last caffeine intake, as in our withdrawal condition, was probably too early to detect changes in sleep structure or subjective sleep quality. In line with this assumption, volunteers subjectively indicated withdrawal symptoms 31 h after caffeine abstinence in the withdrawal condition compared to caffeine. Thus, our findings support the notion that the alterations in sleep structure and subjective sleep quality induced by caffeine abstinence potentially develop at a later stage (> 27 h) of caffeine withdrawal.

Most strikingly and unexpectedly, a reduction in NREM sigma activity during both the withdrawal and caffeine conditions was observed, a phenomenon which is commonly reported under conditions of enhanced sleep pressure.^43-46^ Thus, it seems at first glance in contrast to the reported increases in this frequency range^10,34^ and the well-known alerting effects after acute caffeine intake.^18^ However, during conditions of chronic caffeine intake, mice showed a deeper sleep compared to placebo.^47^ Moreover, repeated caffeine intake enhances the sensitivity of adenosine binding^48^ presumably due to increased adenosine plasma levels,^38^ upregulated adenosine receptors^39-42^ or changes in the functions of adenosine receptor heteromers.^49^ These neuronal alterations in the adenosinergic system might drive the commonly observed changes in the homeostatic sleep-wake regulation such as increased sleepiness when caffeine intake is suddenly ceased.^17^ As reported previously, we also observed in the present study higher subjective sleepiness following caffeine withdrawal when compared to the placebo and caffeine conditions.^25^ Thus, the reduction in sigma activity might reflect adenosinergic changes which already emerge 8 and 15 h after the last caffeine intake in the caffeine and withdrawal condition, respectively. This reduction might reflect withdrawal symptoms which chronic consumers reverse daily by the first caffeine dose. Given the high prevalence of daily caffeine consumers in the society, these findings stress the importance to carefully control for prior caffeine intake when assessing sleep in order to exclude potential confounding by induced withdrawal symptoms which are only detectable in the microstructure of sleep.

Our study has some limitations which must be taken into careful consideration when interpreting the present findings. First, age moderates the effects of caffeine on sleep.^11,14^ Thus, the present results cannot be generalized to other age groups. Second, no genetic information of the study participants is available. A genetic variation of the ADORA2A genotype has been linked with caffeine sensitivity to the effects on sleep.^50^ Thus, carriers of this genetic variance are more likely to curtail caffeine consumption and are consequently excluded from the present study leading to a selection bias. However, the focus of the present study was to investigate habitual caffeine consumers as they represent the majority of the worldwide population.^2^ Third, to reduce variance in the data incurred by the influence of the menstrual cycle on sleep^22^ and the interaction between caffeine metabolism and the use of oral contraceptives,^23,24^ only male volunteers were included which reduces the generalizability of the findings.

In conclusion, we report evidence that daily daytime intake of caffeine and its cessation has no strong effect on sleep structure or subjective sleep quality. However, the quantitative EEG analyses revealed reduced activity in the sigma range during both caffeine and withdrawal. These subtle alterations point to early signs of caffeine withdrawal in the homeostatic aspect of sleep-wake regulation which are already present as early as 8 h after the last caffeine intake. Thus, habitual caffeine consumers constantly expose themselves to a continuous change between presence and absence of the stimulant. Around the clock, their organisms dynamically adapt and react to daily presence and nightly abstinence.

## Acknowledgments

The present work was performed within the framework of a project granted by the Swiss National Science Foundation (320030_163058) and was additionally funded by the Nikolaus und Bertha Burckhardt-Bürgin-Stiftung and the Janggen-Pöhn-Stiftung. Further, we thank our interns Andrea Schumacher, Laura Tincknell, Sven Leach, and all our study helpers for their help in data acquisition and all our volunteers for participating in the study. Moreover, we gratefully acknowledge the help in study organization provided by Dr. Ruta Lasauskaite and the medical screenings conducted by Dr. med. Martin Meyer and Dr. med. Corrado Garbazza.

